# Diverse innate immune factors protect yeast from lethal viral pathogenesis

**DOI:** 10.1101/2022.02.14.480455

**Authors:** Sabrina Chau, Jie Gao, Annette J. Diao, Shi Bo Cao, Amirahmad Azhieh, Alan R. Davidson, Marc D. Meneghini

## Abstract

In recent years, newly characterized anti-viral systems have proven to be remarkably conserved from bacteria to mammals, demonstrating that unique insights into these systems can be gained by studying microbial organisms. Despite the enthusiasm generated by these findings, the key microbial model organism *Saccharomyces cerevisiae* (budding yeast) has been minimally exploited for studies of viral defense, primarily because it is not infected with exogenously transmitted viruses. However, most yeast strains are infected with an endogenous double stranded RNA (dsRNA) virus called L-A, and previous studies identified conserved antiviral systems that attenuate L-A replication. Although these systems do not completely eradicate L-A, we show here that they do prevent proteostatic stress and lethality caused by L-A over-proliferation. Exploiting this new finding, we demonstrate that the genetic screening methods available in yeast can be used to identify additional conserved antiviral systems. Using these approaches, we discovered antiviral functions for the yeast homologs of polyA-binding protein (PABPC1) and the La-domain containing protein Larp1, which are both involved in viral innate immunity in humans. We also identified new antiviral functions for the RNA exonucleases *REX2* and *MYG1*, both of which have distinct but poorly characterized human and bacterial homologs. These findings highlight the potential of yeast as a powerful model system for the discovery and characterization of conserved antiviral systems.

**Significance Statement:** The budding yeast *Saccharomyces cerevisiae* has been minimally exploited for investigation of host-virus interactions despite its chronic infection with a double-stranded RNA virus called L-A. Controverting its presumed harmless nature, we show here that L-A causes pathogenesis in cells lacking parallel-acting viral attenuation pathways. Taking advantage of the genetic tools available in budding yeast, we identify several highly conserved proteins to play a role in antiviral defense. Some of these have been recently identified in humans to be involved in viral innate immunity, thus highlighting the potential of budding yeast as a model organism to identify and investigate new antiviral systems.

## Introduction

All laboratory strains and many environmental isolates of the budding yeast *S. cerevisiae* are infected with a double stranded RNA (dsRNA) virus called L-A (1, 2). L-A belongs to the broadly dispersed *Totiviridae* family of endogenous dsRNA viruses. Like all viruses of this family, the L-A dsRNA genome is packaged within a virion that shields it from host-mediated digestion. Holes in the virion permit the extrusion of RNA transcripts into the cytosol that encode the capsid proteins (Gag) that comprise most of the particle. The L-A transcript also encodes a Gag-pol fusion protein produced at much lower levels than the Gag protein that possesses RNA dependent RNA polymerase activity. Each virion contains a Gag-pol protein, which accounts for L-A replication and transcription within the particle. Encapsidation of the viral transcripts within nascent particles and synthesis of the negative RNA strand by Gag-pol to form the dsRNA genome completes the L-A replication cycle (2). To produce these proteins, L-A employs features typical of RNA viruses found in humans, including a “cap-snatching” mechanism that furnishes L-A transcripts with a 5’-methyl cap and a ribosomal frameshifting mechanism to produce Gag and Gag-pol fusion proteins (3, 4).

Recent studies of bacterial antiviral systems have shown that they share remarkable evolutionary conservation with humans, revealing the potential of microbial organisms for revealing new insights into viral innate immunity (5-11). Indeed, early studies involving L-A led to the discovery of two antiviral systems that have subsequently been shown to contribute to innate immunity against diverse RNA viruses in mammals (12-17). The first of these antiviral systems involves the *SKI2, 3*, and *8* genes, which encode subunits of a conserved ribosome-associated complex that opposes the translation of transcripts that lack poly(A) tails like those encoded by L-A (18-23). A separate pathway of L-A attenuation is provided by Xrn1, a 5’-3’ exoribonuclease that degrades uncapped mRNAs (24-26).

We recently found that the mitochondrial DNA/RNA endonuclease Nuc1 repressed the accumulation of L-A in sporulating cells, representing a new yeast antiviral pathway (27). Nuc1 is a homolog of endonuclease G (EndoG) found in all eukaryotes and many prokaryotes and is most well-known for its role promoting genome fragmentation during mammalian programmed cell death, a prominent last-resort mechanism of viral defense (28, 29). Intriguingly, programmed cell death is intrinsic to yeast sporulation, and Nuc1 fragments the DNA from dying meiotic products during this process in addition to its role in attenuating L-A viral levels that are inherited by the surviving spores (27, 30, 31).

Despite the ubiquitous presence of the L-A virus in lab strains, no fitness consequence has been attributed to it and L-A has earned a reputation as a harmless commensal. Resultingly, although budding yeast has been a powerful system for studies of molecular and cellular biology, it is not widely known as a model for studying antiviral immunity. We show here that L-A infection is deadly for yeast and that it must be actively attenuated through viral innate immunity to preserve viability. Specifically, in strains lacking parallel acting *NUC1* and *SKI* antiviral pathways, L-A copy number is massively increased, and this condition leads to proteostatic stress and lethality at high temperature.

Given the similarities between the L-A viral replication cycle and those of other RNA viruses, we reasoned that further characterization of L-A and the factors that maintain its replication at a low level could reveal new antiviral systems. Identification of conditions that lead to L-A pathogenesis allowed us to use bioinformatic and forward genetic screening approaches to discover new antiviral functions for the RNA exonucleases *REX2* and *MYG1*, both of which have distinct but poorly characterized human and bacterial homologs (32-35). For *MYG1*, we show that its human homolog can perform antiviral activity in yeast. We also identify antiviral functions for the yeast homologs of poly(A)-binding protein (PABPC1) and the La-domain containing protein Larp1, which are both involved in viral innate immunity in humans (36, 37). These findings provide new examples of innate immune conservation from microbes to humans and highlight the potential of yeast as a powerful model system for the discovery of new antiviral mechanisms.

## Results

### The NUC1, SKI and XRN1 yeast antiviral systems collaborate to prevent L-A pathogenesis

Our previous studies of *NUC1* were focused on meiotic cells (27). To investigate *NUC1* antiviral function in vegetatively growing yeast, we examined L-A copy number in mitotic haploid cells in the reference BY4742 strain background. We observed the levels of L-A dsRNA using ethidium bromide staining of electrophoresed RNA and found that while *nuc1*Δ and *ski3*Δ single mutants had levels similar to wild type, a *nuc1*Δ *ski3*Δ double mutant showed a large increase in L-A dsRNA **(Fig. 1A)**. We corroborated these findings using immunofluorescence microscopy with a dsRNA antibody widely used to detect replicating RNA viruses (38, 39). These images showed that L-A dsRNA accumulated in foci, reminiscent of “viral factory” sites of viral replication observed in human cells **(Fig. 1B and S1)** (40). Consistent with previous findings in other strain backgrounds (24, 27, 41), western blotting showed that Gag protein levels were elevated in the *nuc1*Δ and *ski3*Δ mutants **(Fig. 2A)**. Furthermore, we showed that a *nuc1*Δ *ski3*Δ double mutant accumulated a massively elevated level of Gag **(Fig. 2A)**. These data show that *NUC1* and *SKI3* participate in separate antiviral pathways and that loss of both pathways results in a greatly increased L-A viral load.

**Figure 1.**
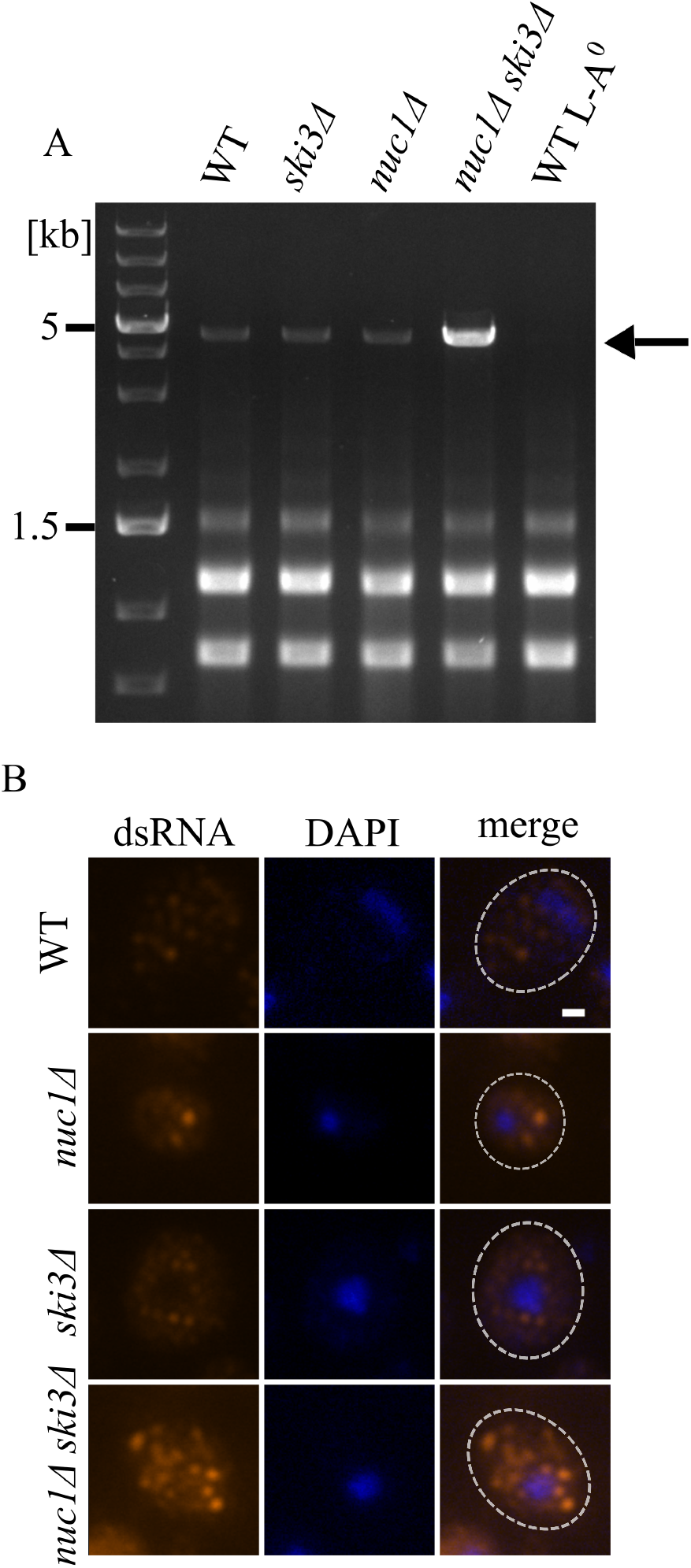
Loss of *NUC1* and *SKI3* displays a synergistic increase in L-A viral load. **(A)** An ethidium bromide-stained gel of total RNA prepared from the indicated strains is shown, with the 4.6 Kb L-A dsRNA band indicated with an arrow. **(B)** Immunofluorescence was used to visualize L-A dsRNA (red) in cells of the indicated genotypes. These strains were cured of the weakly abundant L-BC dsRNA virus to eliminate background staining (see Methods). DAPI staining of DNA is shown in blue. Scale bar, 1 µm.

**Figure 2.**
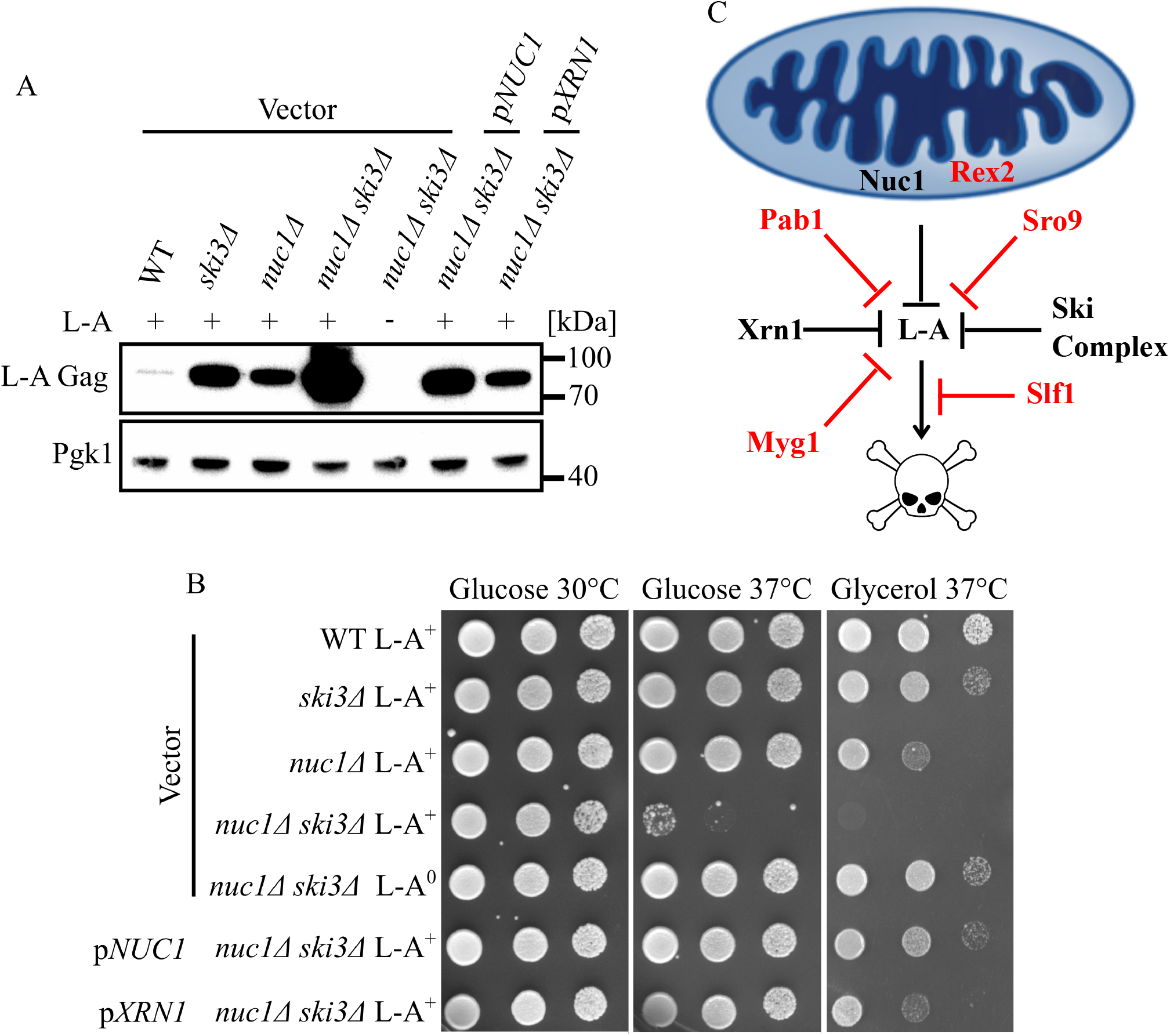
L-A attenuation protects yeast from lethal pathogenesis. **(A)** Western blotting of L-A Gag and Pgk1 protein levels in the indicated strains is shown. Molecular weight markers are indicated on the right. **(B)** Spot test growth assays of the strains from 2A is shown. Strains were spotted on -Leu media containing either glucose or glycerol and grown at the indicated temperatures. **(C)** The mitochondrial proteins Nuc1 and Rex2 collaborate with the cytosolic proteins Xrn1, SkiC, and Myg1 to regulate L-A protein level and ensure cell fitness. The translation control factors Pab1 and Sro9 alleviate L-A pathogenesis by suppressing L-A level while Slf1, the paralog of Sro9, suppress L-A pathogenesis without affecting L-A level. Newly discovered L-A attenuation factors are indicated in red.

To determine if high L-A viral load affects cell fitness, we examined cellular growth using spot test growth assays. Subtle growth defects of *nuc1*Δ and *ski3*Δ single mutants were observed at 37° C when cells were grown with glycerol rather than glucose as the carbon source, a condition under which yeast relies on mitochondrial respiration **(Fig. 2B)**. Remarkably, although *nuc1*Δ *ski3*Δ double mutants grew normally at 30° C, they exhibited conditional lethality at 37° C regardless of carbon source **(Fig. 2B)**. As expected, viability at high temperature was restored to a *nuc1*Δ *ski3*Δ double mutant by a *NUC1*-expressing plasmid which elicited a corresponding decrease in Gag levels (**Fig. 2A and 2B**). To confirm that the growth defect of the *nuc1*Δ *ski3*Δ double mutant was caused by L-A, we constructed an isogenic strain cured of L-A (L-A^0^) and assayed for its growth at high temperature. We found that the growth defect was completely alleviated, implying that the conditional lethality was a result of unrestricted L-A replication **(Fig. 2B)**.

To further test how *NUC1* interacts with known antiviral pathways, we characterized its relationship with *XRN1*. We found that a *nuc1*Δ *xrn1*Δ double mutant accumulated greatly elevated levels of Gag compared with either single mutant and exhibited L-A-dependent conditional lethality at high temperature **(Fig. S2A and S2B)**, suggesting that *NUC1* and *XRN1* act in parallel pathways to attenuate L-A. Reflecting their key and non-redundant roles in bulk mRNA regulation, an *xrn1*Δ *ski3*Δ double mutant is inviable, even in strains lacking L-A (42). To determine if *XRN1* represents an antiviral system independent of both *NUC1* and *SKI3*, we used a high-copy plasmid to overexpress *XRN1* in a *nuc1*Δ *ski3*Δ double mutant. Indeed, we observed a substantial decrease in Gag levels and suppression of the *nuc1*Δ *ski3*Δ conditional lethality using plasmid driven *XRN1* over-expression **(Fig. 2A and 2B)**. We conclude that Nuc1, Ski3 and Xrn1 convergently oppose L-A replication and that massively increased L-A viral load in *nuc1*Δ *ski3*Δ mutants caused lethal pathogenesis at high temperature **(Fig. 2C)**.

### A bioinformatic-based genetic screen for new viral attenuation fatctors

The L-A-dependent conditional lethality of *nuc1*Δ *ski3*Δ double mutants opened the possibility that other antiviral factors could be identified through combinatorial mutant studies. To identify new candidate antiviral factors, we searched a curated genetic interaction database for gene deletions that caused a synthetic growth defect when combined with *nuc1Δ* in at least two high-throughput screening studies (43). In addition to the expected presence of *XRN1* and *SKI* deletions in this data set, we found sixteen additional genes. We used genetic crossing to make triple mutants combining deletions of each of these sixteen genes with *nuc1*Δ *ski3*Δ and found five that caused severe L-A dependent growth defects (**Table S1**). Here we describe two of these genes that we have confirmed as new antiviral factors.

The first new antiviral gene identified in our screen, *REX2*, encodes a 3’-5’ RNA exonuclease conserved from bacteria to humans (33). Both Rex2 and its human homolog REXO2 localize to the mitochondria and contain an EXOIII domain widely found in prokaryotic and eukaryotic proteins, including the interferon stimulated antiviral protein ISG20 (34, 35, 44-46). We found that a *rex2*Δ *nuc1*Δ double mutant strain accumulated greatly increased levels of Gag and exhibited L-A-dependent lethality at high temperature **(Fig. 3A and 3B)**. Remarkably, *nuc1*Δ *ski3*Δ *rex2*Δ and *nuc1*Δ *xrn1*Δ *rex2*Δ triple mutants were inviable under all growth conditions and these defects were reversed in L-A^0^ strains **(Fig. S3)**. These findings demonstrate the severe pathogenic potential of the L-A virus and identify a new antiviral role for a highly conserved mitochondrially localized RNA exonuclease **(Fig. 2C)**.

**Figure 3.**
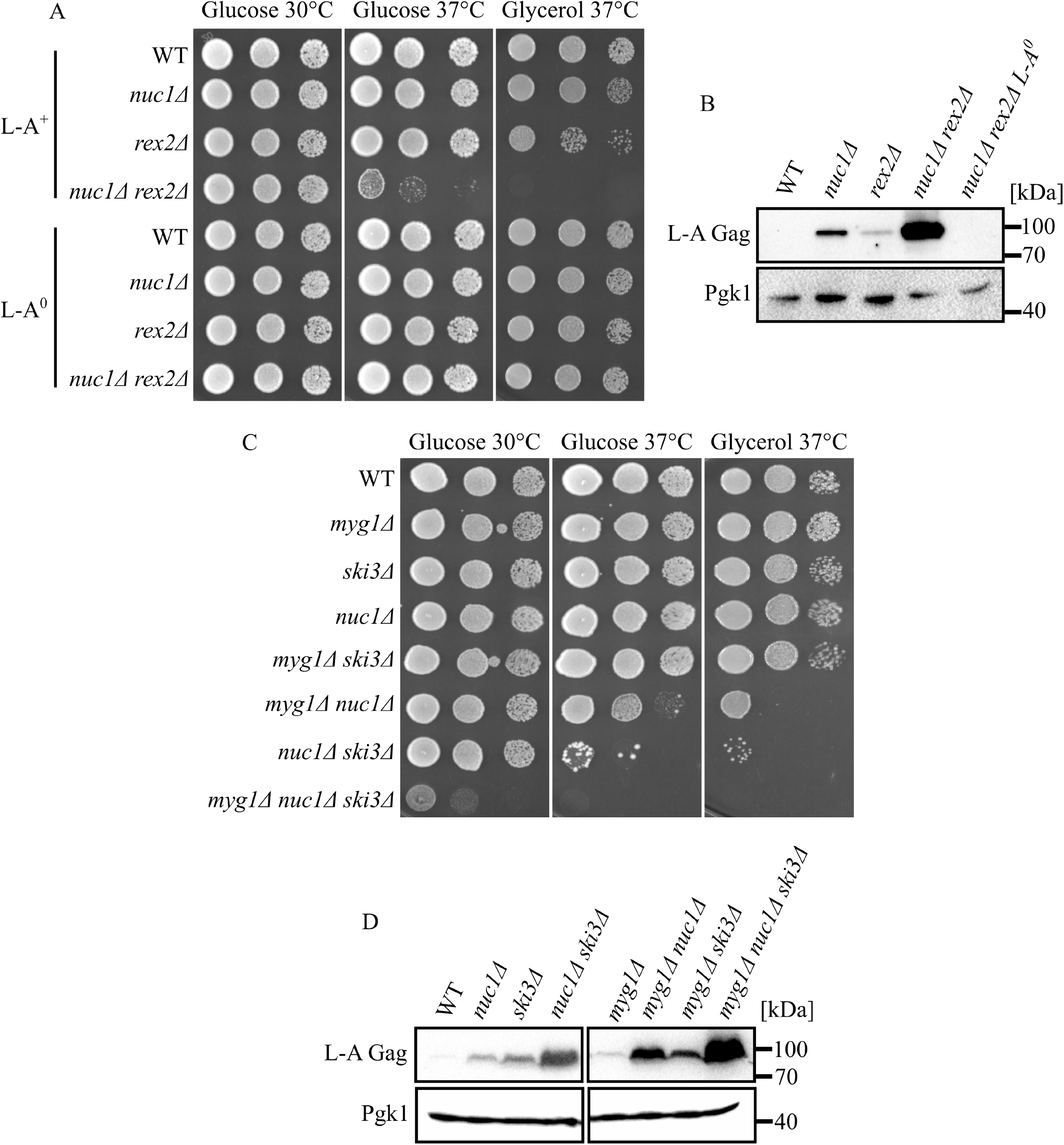
New antiviral factors are identified by exploiting L-A pathogenesis. **(A)** Spot analysis of strains defective in *NUC1* and *REX2* is shown. Strains are spotted on SC media containing either glucose or glycerol and grown at the indicated temperature. **(B)** Western blotting of L-A Gag and Pgk1 protein levels of strains in Figure 3A. Molecular weight markers are indicated on the right. **(C)** Spot analysis of strains defective in *MYG1, NUC1*, and *SKI3* is shown. Strains are spotted on SC media containing either glucose or glycerol and grown at the indicated temperature. **(D)** Western blotting detection of L-A Gag and Pgk1 protein levels in the indicated strains is shown. Molecular weight markers are indicated on the right.

The second gene identified in our screen was *MYG1*, the yeast homolog of human **<u>M</u>**elanoc**<u>Y</u>**te proliferation **<u>G</u>**ene **<u>1</u>**, a 3’-5’ RNA exonuclease which has homologs found in all taxa (32). A *myg1*Δ *nuc1*Δ double mutant strain exhibited a large increase of Gag accumulation compared to the single mutants and displayed a severe L-A-dependent growth defect at high temperature **(Fig. 3C, 3D, and S4A)**. We were able to recover *nuc1*Δ *ski3*Δ *myg1*Δ triple mutants, though they were extremely slow growing at 30° C and accumulated even higher levels of Gag **(Fig. 3C and 3D)**. These growth defects were also reversed in L-A^0^ strains **(Fig. S4A)**. *MYG1* thus represents a new viral attenuation factor, acting in parallel to both *NUC1* and the *SKI* complex **(Fig. 2C)**.

Mutations leading to overexpression of human MYG1 are associated with the auto-immune disorder vitiligo, suggesting that human MYG1 may play some role in normal innate immunity (47, 48). We explored this possibility using a plasmid expressing human MYG1 under the control of a constitutive yeast promoter (32) and found that human MYG1 rescued the conditional growth defect of a *nuc1*Δ *myg1*Δ mutant **(Fig. S4B)**. These findings show that the antiviral function of yeast *MYG1* can be accomplished by human MYG1.

### High copy suppression screening identifies new yeast antiviral factors that are conserved in humans

Since *XRN1* over-expression suppressed the growth defects of a *nuc1*Δ *ski3*Δ strain, we hypothesized that over-expression of other antiviral factors would produce a similar effect and that this phenomenon could be used as a screen to identify new antiviral systems. We employed a high-copy plasmid suppression screen to detect genes whose overexpression alleviated the conditional lethality of a *nuc1Δ ski3Δ* strain (see Methods). Using this screen, we identified *SRO9, SLF1*, and *PAB1* as high-copy suppressors of *nuc1*Δ *ski3*Δ, all of which encode ribosome-associated RNA binding proteins **(Fig. 4A)** (49, 50). Sro9 and Slf1 are paralogous **<u>l</u>**upus-**<u>a</u>**utoantigen (La) domain containing proteins broadly found in eukaryotes. Notably, their human homolog, Larp1, was recently identified in screens for proteins bound to the SARS-Cov-2 plus strand ssRNA or nucleocapsid (36, 51). Larp1 was a major focus in one of these studies and was shown to attenuate SARS-Cov-2 replication in human cells, though its mechanism is not known (36). *PAB1* encodes the highly conserved <u>P</u>oly<u>A</u>-<u>B</u>inding <u>P</u>rotein, which is a common target of viral inhibition in humans through diverse mechanisms (37).

**Figure 4.**
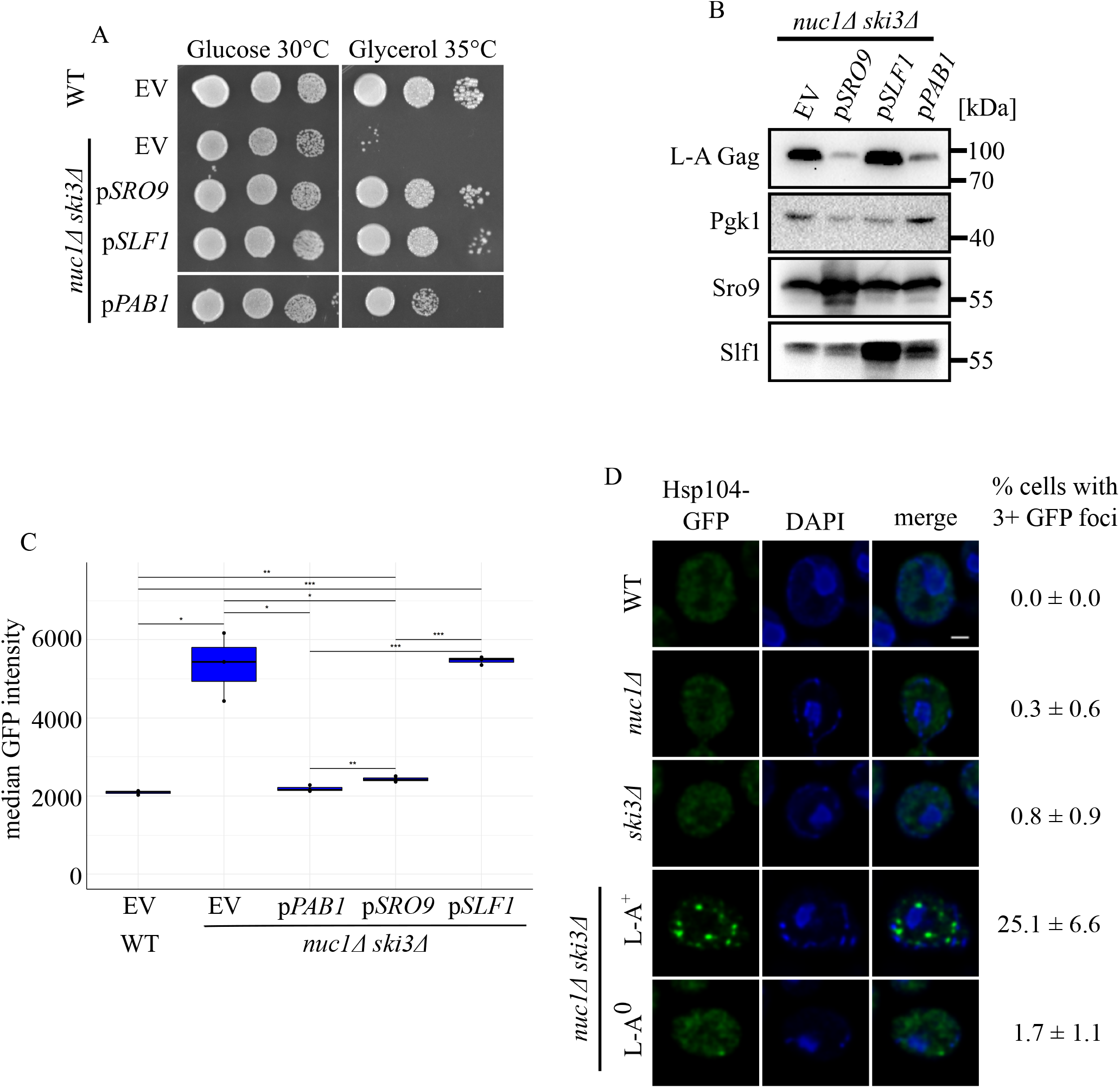
Overexpression of translation control factors alleviate L-A pathogenesis. **(A)** Spot test growth assays of the high copy suppressors *SRO9, SLF1*, and *PAB1* are shown. **(B)** Western blotting detection of L-A Gag, Pgk1, Sro9, and Slf1 protein levels in the strains from 4A is shown. Molecular weight markers are indicated on the right. **(C)** Flow cytometry was used to measure HSE-GFP expression in the indicated strains. Median GFP intensity is shown. n = 3. * *p* < 0.05, ** *p* < 0.01, *** *p* < 0.001. The *p* value was calculated using the unpaired student’s t-test. **(D)** Fluorescence microscopy of Hsp104-GFP in indicated strains. Cells were stained with DAPI to visualize nuclei. Scale bar, 1 µm. Percentage of cells with 3+ GFP foci is shown on the right. n = 3. 75-140 cells are counted for each replicate.

We found that overexpression of *PAB1* or *SRO9* markedly reduced Gag levels in a *nuc1*Δ *ski3*Δ mutant, explaining their rescuing phenotypes **(Fig. 4B)**. Curiously, even though *SLF1* overexpression rescued the *nuc1*Δ *ski3*Δ growth defect just as well as *SRO9*, it did not lead to any reduction in Gag levels **(Fig. 4A and 4B)**. These findings suggest that *PAB1* and *SRO9* rescue cells by suppressing L-A replication and that *SLF1* protects cells from the pathogenic consequences of elevated viral replication **(Fig. 2C)**.

### L-A pathogenesis is associated with proteostatic stress

To gain insight into the divergent mechanisms of Sro9 and Slf1 antiviral activities, we considered what the physiological consequences L-A pathogenesis could be and how *SRO9*/*SLF1* might differentially impact them. We noted a previous study in which deletions of *NUC1* or of *SKI*-complex genes led to weak induction of a GFP reporter gene controlled by Hsf1 (52), a conserved transcription factor that senses proteostatic stress and activates the gene expression response (53-55). Using flow cytometry with this reporter (HSE-GFP), we confirmed these results and determined that a *nuc1*Δ *ski3*Δ double mutant caused synergistic and L-A-dependent activation of HSE-GFP **(Fig. 4C and S5A)**. We hypothesized that the massive accumulation of Gag observed in *nuc1*Δ *ski3*Δ mutants accounted for this proteostatic stress response. Supporting this, HSE-GFP activation of a *nuc1*Δ *ski3*Δ double mutant was reverted by *PAB1* or *SRO9* over-expression, mirroring these genes consequences for Gag accumulation **(Fig. 4B and 4C)**. Surprisingly, over-expression of the *SRO9* paralog *SLF1* did not prevent HSE-GFP activation. The evolutionary divergence of *SRO9* and *SLF1* has thus resulted in different antiviral mechanisms, with *SRO9* suppressing viral protein accumulation and associated proteostatic stress, and *SLF1* possibly protecting cells from the toxic consequences of viral induced proteostatic stress **(Fig. 2C)**.

Proteostatic defects are often associated with the accumulation of cytotoxic protein aggregates that can be visualized using a GFP fusion of Hsp104, a protein disaggregase that localizes to sites of protein aggregation (56). To further explore the proteostatic defects associated with L-A pathogenesis we used fluorescence microscopy to visualize Hsp104-GFP foci in a variety of strains. As expected, wild type cells grown at 30° C rarely accumulated observable Hsp104-GFP foci. While *nuc1*Δ and *ski3*Δ single mutants resembled wild type, strikingly, a *nuc1*Δ *ski3*Δ double mutant exhibited more than 25% of cells with 3 or more Hsp104-GFP foci **(Fig. 4D)**. As with all other phenotypes we have observed for *nuc1*Δ *ski3*Δ, its accumulation of Hsp104-GFP foci was dependent on the presence of L-A **(Fig. 4D)**. These findings show that high viral load caused by deletion of *NUC1* and *SKI3* led to the accumulation of Hsp104-GFP foci indicative of cytotoxic protein aggregation.

## Discussion

The L-A dsRNA virus infects all lab strains and accompanies virtually every yeast experiment. Despite its ubiquitous presence, L-A has been largely overlooked due to its apparent benign nature. Here, we show that L-A has profound consequences for yeast when its replication is uncontrolled, and that diverse innate immune pathways maintain L-A replication at a tolerable level. Specifically, we show that in strains lacking the parallel-acting *NUC1* and *SKI3* antiviral genes, L-A replication is massively upregulated, leading to proteostatic stress and conditional lethality at high temperature. Leveraging this new discovery, we used bioinformatic and forward genetic screens to identify new yeast genes that function to restrict L-A replication or to protect cells from the pathogenic consequences of unrestrained L-A replication. As these screens were not saturating, they suggest that the yeast genome likely encodes numerous other antiviral/viral protective factors.

Given the clear risk of L-A infection, how it nevertheless persists in the face of ever-present attenuation is puzzling. An explanation for this paradox is that L-A provides a counterbalancing benefit. One possible benefit of L-A is that it enables some strains to maintain satellite viruses that encode secreted toxins that kill neighboring uninfected cells. However, L-A is present in many strains that lack “Killer” satellites, so this explanation is insufficient to explain the persistence of L-A infection. We thus speculate that L-A may have some additional benefit that counterbalances its deleterious potential.

Our discovery of Rex2 as a viral attenuation factor expands the arsenal of known mitochondrial antiviral factors beyond Nuc1 and suggests that mitochondria are a key antiviral hub in yeast. Indeed, mitochondria serve central roles in viral defense as a programmed cell death regulator and as a platform for antiviral signaling in humans. How do mitochondrial nucleases attenuate a virus that resides in the cytosol in yeast? One possibility is that these enzymes, while targeted to mitochondria, may nevertheless accumulate to low but sufficient levels in the cytosol to accomplish L-A attenuation directly. Consistent with this hypothesis, we showed previously that Nuc1 accumulates to low levels in the cytosol of meiotic cells, though our methods could not detect it in the cytosol of mitotic cells (27). Another hypothesis is that some aspect of the L-A replication cycle occurs in intimate association with mitochondria. For example, L-A transcripts may associate with and possibly traverse mitochondria, thus exposing them to Nuc1 and/or Rex2. It is interesting to note that human MYG1 protein exhibits dual mitochondrial/nucleolar localization, though yeast Myg1 has only been observed in the nucleus and cytosol (32). Our results highlight a potential general importance of mitochondria for viral innate immunity in eukaryotes and position the yeast-L-A system as a powerful model for further studies of this topic.

The antiviral *SKI* complex associates with translating ribosomes and our identification of Pab1, Sro9, as Slf1 as high copy suppressors of L-A pathogenesis further reveal the translating ribosome as a key hub of yeast antiviral activity. The finding that *PAB1* (polyA binding protein) repressed L-A is surprising given the absence of polyA tails in L-A transcripts, suggesting that Pab1 does not act on L-A directly. Previous findings showed that L-A transcripts compete with polyA+ yeast mRNAs for capture of 60S ribosomal subunits to form translating 80S complexes (57). One model explaining these findings is that Pab1 enhances the translation of polyA tail containing mRNAs, which then deplete the availability of 60S subunits for L-A transcripts for translation. The roles of Sro9 and Slf1 in translation n are less well understood, but their functions may similarly relate to competition of L-A transcripts for 60S subunits. Importantly, the homologs of Pab1 and Sro9/Slf1 are involved in human viral defense and further studies of these genes in yeast will shed light on antiviral mechanisms conserved from yeast to human.

## Method Details

### Strains, media, and plasmids

Standard *S. cerevisiae* genetic and strain manipulation techniques were used for strain construction. Strains were grown at 30°C in either YPAD or synthetic complete media with appropriate amino acid drop out for plasmid maintenance unless otherwise specified. Refer to Table **S2** for strains and plasmids used in this paper. The yOB255 *URA3::HSE-EmGFP* strain was obtained from Onn Brandman(58). The p5476 *2μ LEU2*, p5476 *NUC1 2μ LEU2*, p5476 *XRN1 2μ LEU2*, p5476 *PAB1 2μ LEU2*, p5476 *SRO9 2μ LEU2*, p5476 *SLF1 2μ LEU2* MOBY plasmids were obtained from Brenda Andrews (59). The pVB3011 *L-BC GAG (P*_*GAL1*_) plasmid was obtained from Suzanne Sandmeyer (60). The RLY8470 *trp::mCherry-FIS1TM-KanMX; HSP104-GFP-HIS3* strain was obtained from Rong Li (61).

### Spot analysis

Yeast strains were grown overnight at 30°C to saturation. Each strain was diluted to an OD_600_ = 0.4, serial diluted four times by 10-fold, and spotted onto agar plates containing synthetic complete media or drop out media supplemented with the indicated carbon source (2% glucose or 3% glycerol).

### Protein extraction and Western blot

5 OD of log-phase cells were harvested and permeabilized with 0.1N NaOH at room temperature for 5 minutes. The cells were then pelleted and resuspended in SDS/PAGE buffer before heating at 100°C for 10 minutes. The samples were centrifuged to isolate the soluble fraction for Western blotting. Protein concentrations were determined with an RC/DC assay (BioRad 5000121). Equal amounts of protein were electrophoresed on 10% SDS/PAGE and transferred to poly(vinylidene difluoride) membranes. Blots were incubated in primary antibody overnight and probed with 1:3,000 horseradish peroxidase (HRP)-conjugated horse anti-mouse (7075; Cell Signaling Technology) or goat anti-rabbit (7074; Cell Signaling Technology) secondary antibody. The proteins were detected with Luminata Forte Western HRP Substrate (EMD Millipore) or SuperSignal™ West Atto Ultimate Sensitivity Substrate (A38555, Thermo Scientific). Blots were imaged with the Bio-Rad ChemiDoc XRS+ system, and images were processed with the Image Lab software package (Bio-Rad). The primary antibodies and their dilutions were 1:5,000 anti-Pgk1 (ab113687; Abcam), 1:2,000 anti-L-A Gag (obtained from Reed Wickner), 1:8000 anti-Hsp60 (received from Toshiya Endo)(62), 1:500 anti-Tom22(62), 1:2000 anti-Om45(62), 1:1000 anti-Cox2 (ab110271; Abcam), 1:1000 anti-Sro9, and 1:1000 anti-Slf1 (received from Sandra Wolin)(49).

### Virus curing

L-A^0^ strains were generated similar to as we described previously(27). Briefly, a progenitor L-A^0^ strain was isolated following treatment with 32 mM Anisomycin (BioShip ANS245) for 4 days in YPD media and plating for single colonies. Isolates were tested for the presence of the L-A genome by RT-PCR. This L-A^0^ strain was then used for subsequent backcrossing with *mak3Δ* to generate other L-A^0^ strains.

To evict L-BC, cells were transformed with the *L-BC GAG (P*_*GAL1*_) plasmid and grown overnight in – URA drop out media supplemented with 2% raffinose. The saturated culture was diluted and grown in raffinose to log phase before treated with 2% galactose for 24 hours. The resulting culture was plated for single colonies on an agar plate containing synthetic complete media supplemented with 5-FOA. Isolates were tested for the presence of the L-BC genome by RT-PCR.

### RNA extraction and reverse transcription

10 OD of log phase cells was harvested and resuspended in phenol solution (P4682; Sigma-Aldrich), SDS, and buffer AE (10 mM Tris·Cl and 0.5 mM EDTA, pH 9.0). Samples were incubated at 65°C for 30 minutes, and the phase-separated supernatant was washed with chloroform and precipitated in isopropanol overnight. The precipitate was washed with 70% ethanol and dissolved in water. RNA samples were then purified with the RNeasy Mini Kit (Qiagen). Four micrograms of RNA were reverse-transcribed using random nonamers and Maxima H Minus Reverse Transcriptase (Thermo Fisher). The cDNA product was isolated by alkaline hydrolysis and treated with RNase A.

### Immunofluorescence

Log phase cells were fixed with 4% formaldehyde and washed 2 times with 1x phosphate-buffered saline (PBS). Cells were resuspended in sorbitol buffer (1.2 M sorbitol and 20 mM potassium phosphate buffer, pH 7.2) and spheroplated with β-mercaptoethanol (M7522; Sigma-Aldrich) and zymolase at 30°C. Spheroplasted cells were added onto polylysine coated slides and incubated with primary antibody 1:1000 J2 anti-dsRNA (SCICONS) overnight. The cells were detected with 1:1000 cyanine3 conjugated goat anti-mouse (Jackson ImmunoResearch Laboratories) and stained with DAPI. The images were acquired on the Zeiss Axio Imager Z1 with Volocity and processed with ImageJ.

### Flow cytometry

Log phase cells were stained with propidium iodide, fixed with 4% formaldehyde, and washed 3 times with 1x PBS. Fluorescence intensity was measured with Becton Dickinson LSR II flow cytometer and analyzed with Flowing Software. The median GFP intensities of three biological replicates were plotted using R studio ggplot2.

### Fluorescence microscopy

Log phase cells were fixed with 4% formaldehyde and washed 3 times with 1x PBS. Fixed cells were mounted onto microscopy slide and stained with DAPI. Images were acquired with Leica Sp8 confocal LSM and processed with Leica Application Suite (LAS X). The deconvolution program Lightning was applied to all images.

### High copy suppression screen

A plasmid library of random genomic inserts in the YEP24 plasmid backbone was transformed into a *nuc1Δ ski3Δ* strain and grown on -URA agar plates at high density (63). Lawns of transformants grown at 30° C were replica plated to 37° C and potential suppressors were obtained from the colonies that grew at this temperature. To confirm that the suppression was plasmid dependent, the potential suppressors were first grown on agar plate containing 0.1% 5-fluoroorotic acid (5-FOA) for counterselection and then transferred to a fresh agar plate at 37°C supplemented with glycerol. The plasmids of the candidates were isolated and sequenced to identify potential genes responsible for the suppression of L-A pathogenesis when over-expressed. The candidate genes were confirmed by spot analysis using p5476 MOBY plasmids.

### Quantification and Statistical Analysis

Data was tested for statistical significance using Excel and plotted with R studio ggplot2. The details of the statistical tests are described in the figure legends.

## Acknowledgements

We are grateful Dr. Charlie Boone and Dr. Brenda Andrews for generous sharing of strains and plasmid and to Angus McQuibban for critical comments on the manuscript. M.D.M. is funded by NSERC RGPIN-2019-07230. A.R.D. is funded by CIHR FDN-15427.

## Author Contributions

S.C., J.G., A.D., S.B.C., and M.D.M. conceived, planned, and executed the experiments.

S.C., J.G., A.D., and M.D.M. analyzed the data. S.C., A.R.D., and M.D.M. wrote the manuscript.

## Data and Material Availability

All data is available in the main text or supplementary materials.

---

**Figure S1.**
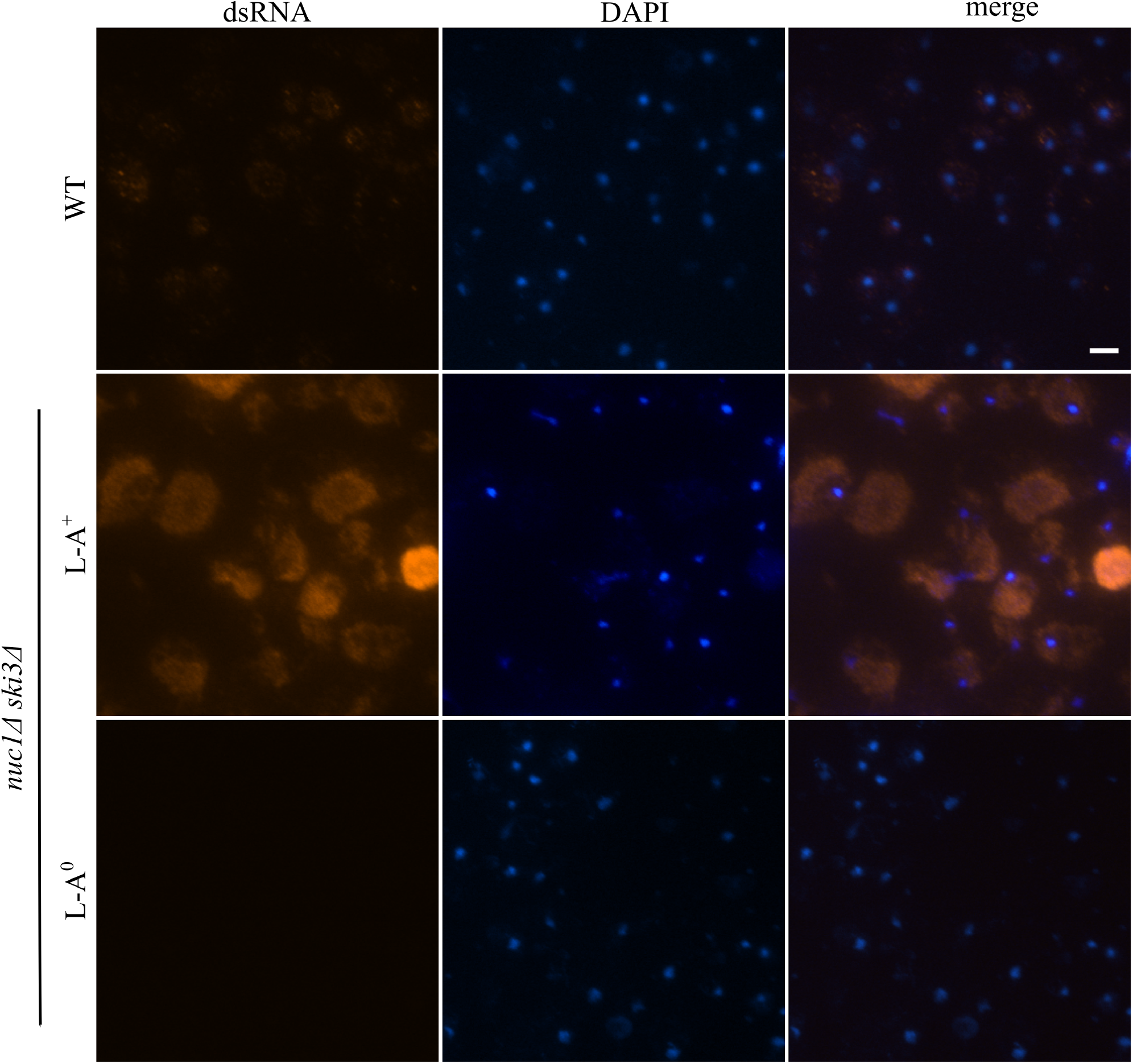
Additional images of immunofluorescence of dsRNA (red) of strains from Figure 1B. Images of fields of cells of indicated genotype is shown. These strains were cured of the weakly abundant L-BC dsRNA virus to eliminate background staining (see Methods). Cells are stained with DAPI to visualize nuclei. Scale bar, 30 µm.

**Figure S2.**
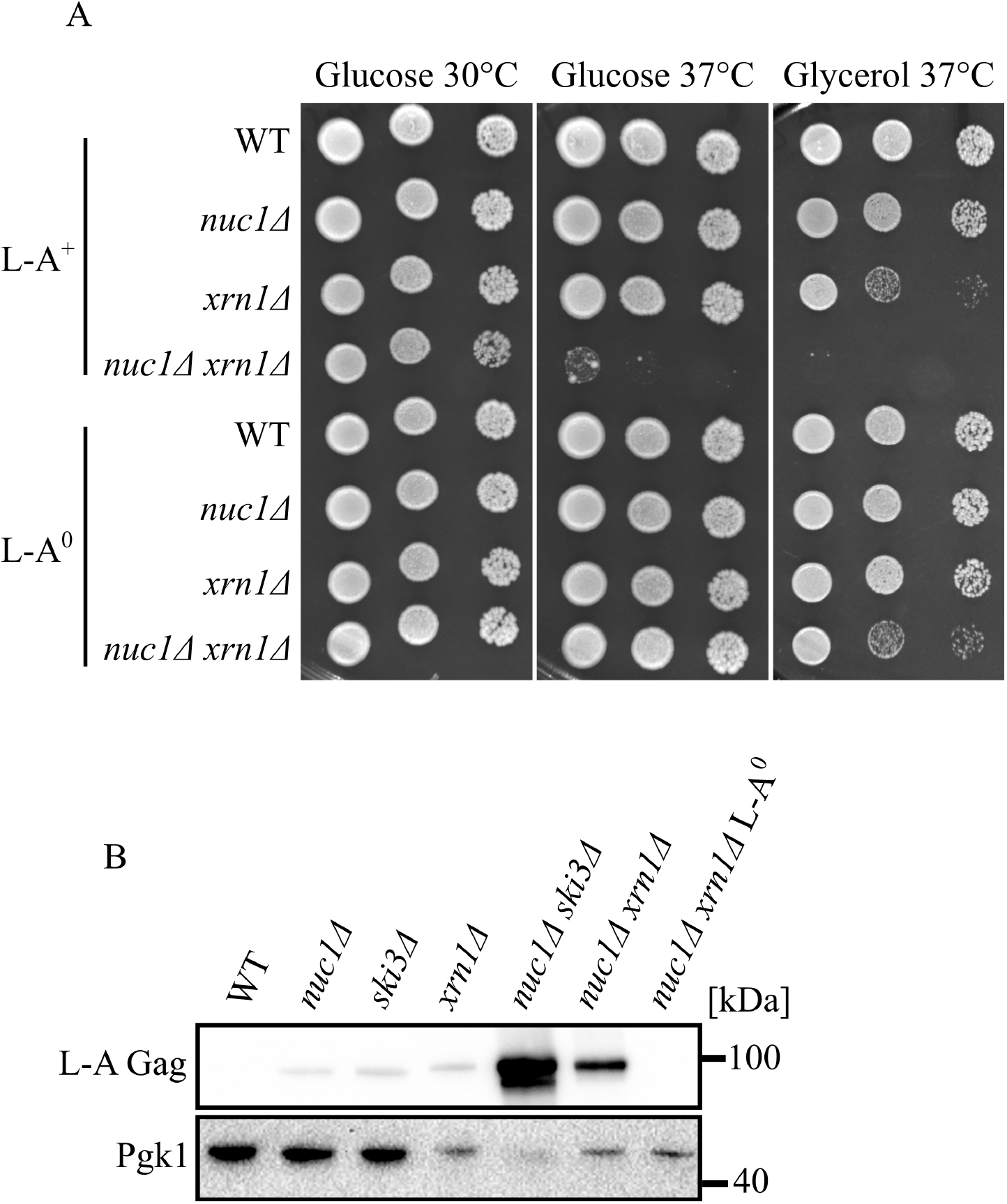
*XRN1, NUC1*, and *SKI3* work in parallel pathways to regulate L-A Gag level. **(A)** Spot analysis of strains defective in *NUC1* and *XRN1* is shown. Strains are spotted on SC media containing either glucose or glycerol and grown at the indicated temperature. **(B)** Western blotting of L-A Gag and Pgk1 protein levels of strains in Figure S2A. Molecular weight markers are indicated on the right.

**Figure S3.**
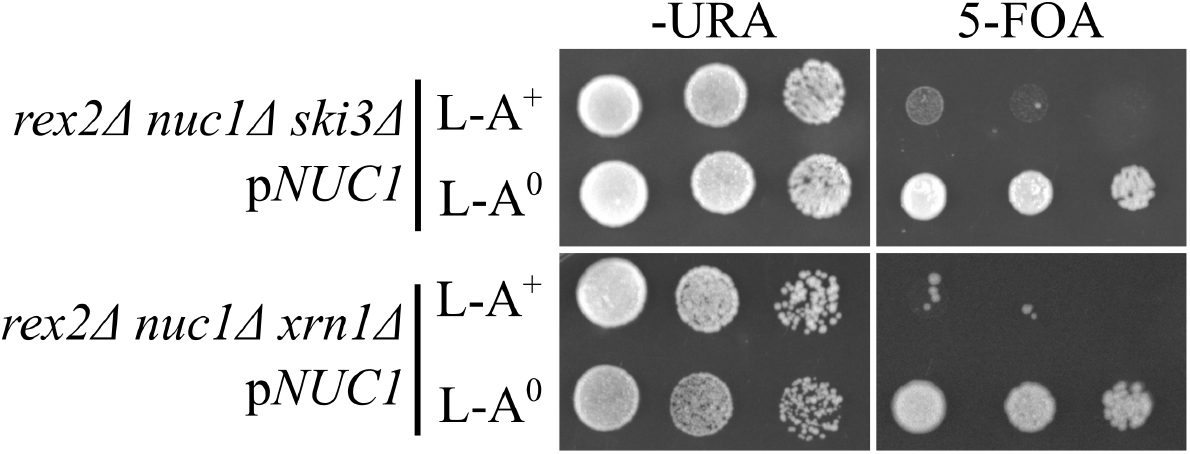
The mitochondrial exonuclease *REX2* is an antiviral factor. Spot analysis of strains defective in 3 parallel antiviral pathways containing plasmid expressing *NUC1* is shown. Strains are spotted on -URA media or synthetic complete (SC) media supplemented with 0.1% 5-fluoroorotic acid (5-FOA).

**Figure S4.**
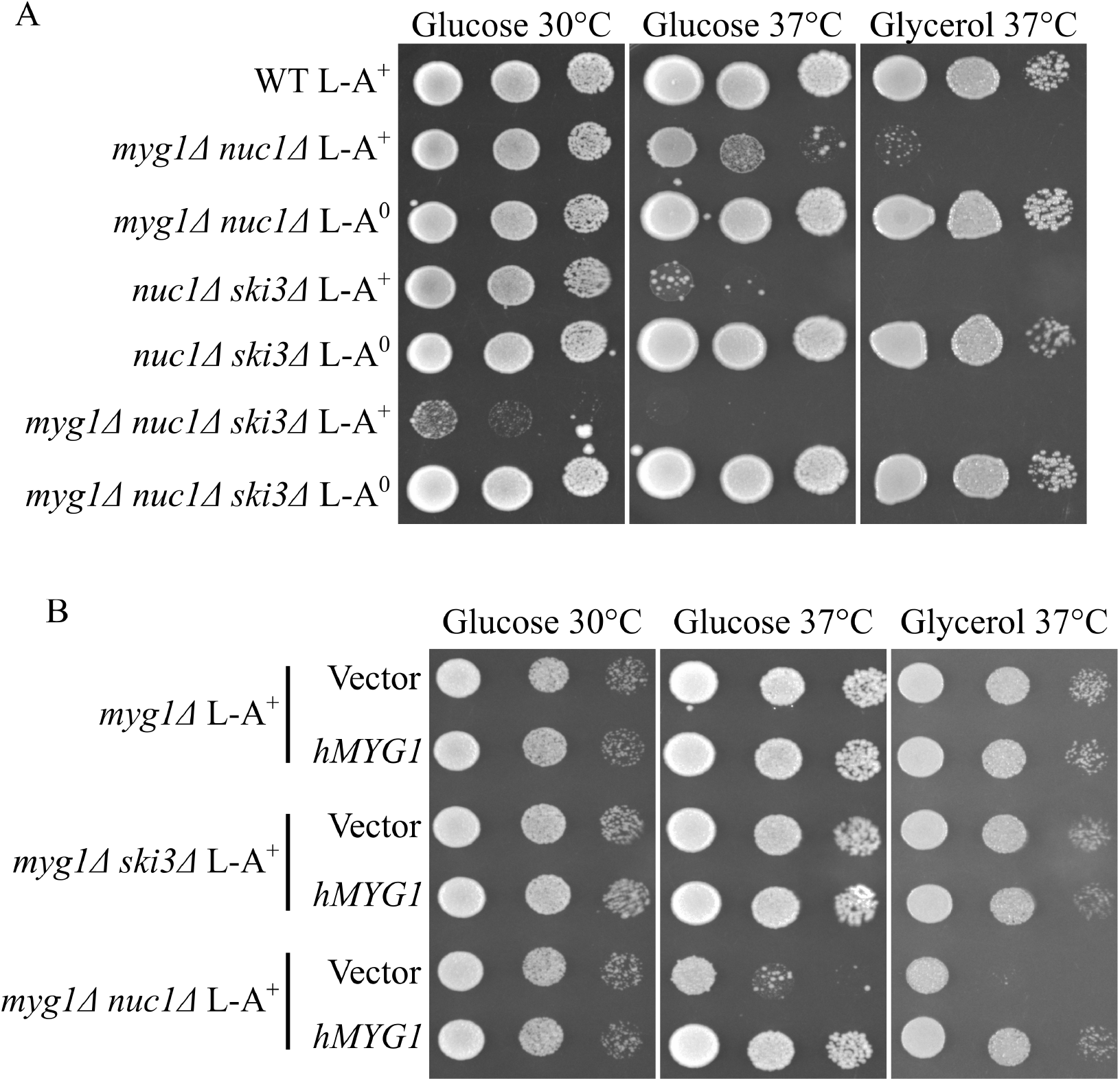
*MYG1* collaborates with *NUC1* and *SKI3* to ensure cell fitness in a L-A dependent manner. **(A)** Spot analysis of strains defective in *MYG1, NUC1*, and *SKI3* with and without L-A is shown. Strains are spotted on SC media containing either glucose or glycerol and grown at the indicated temperature. **(B)** Spot analysis of strains from S4A expressing *hMYG1* on a plasmid. Strains are spotted on -URA media containing either glucose or glycerol and grown at the indicated temperatures.

**Figure S5.**
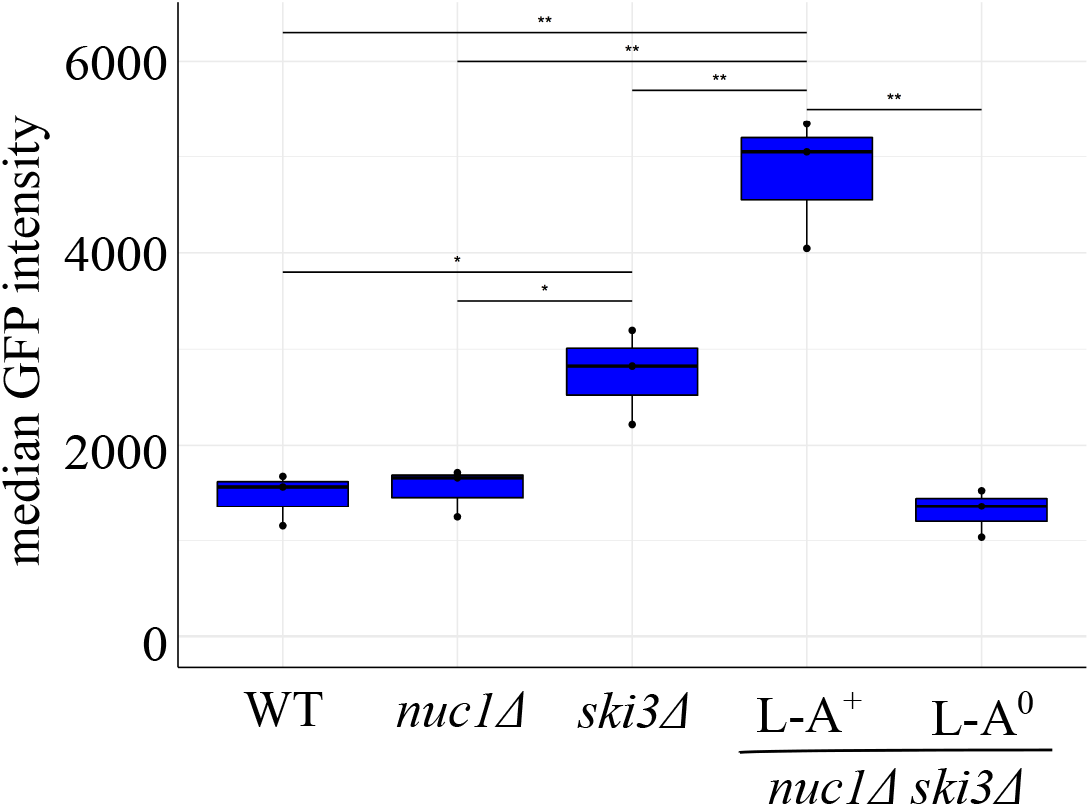
L-A pathogenesis correlates with proteostatic stress. Flow cytometry was used to measure HSE-GFP expression in the indicated strains. Median GFP intensity is shown. n = 3. * *p* < 0.05, ** *p* < 0.01. The *p* value was calculated using unpaired student’s t-test.

**Table S2.**
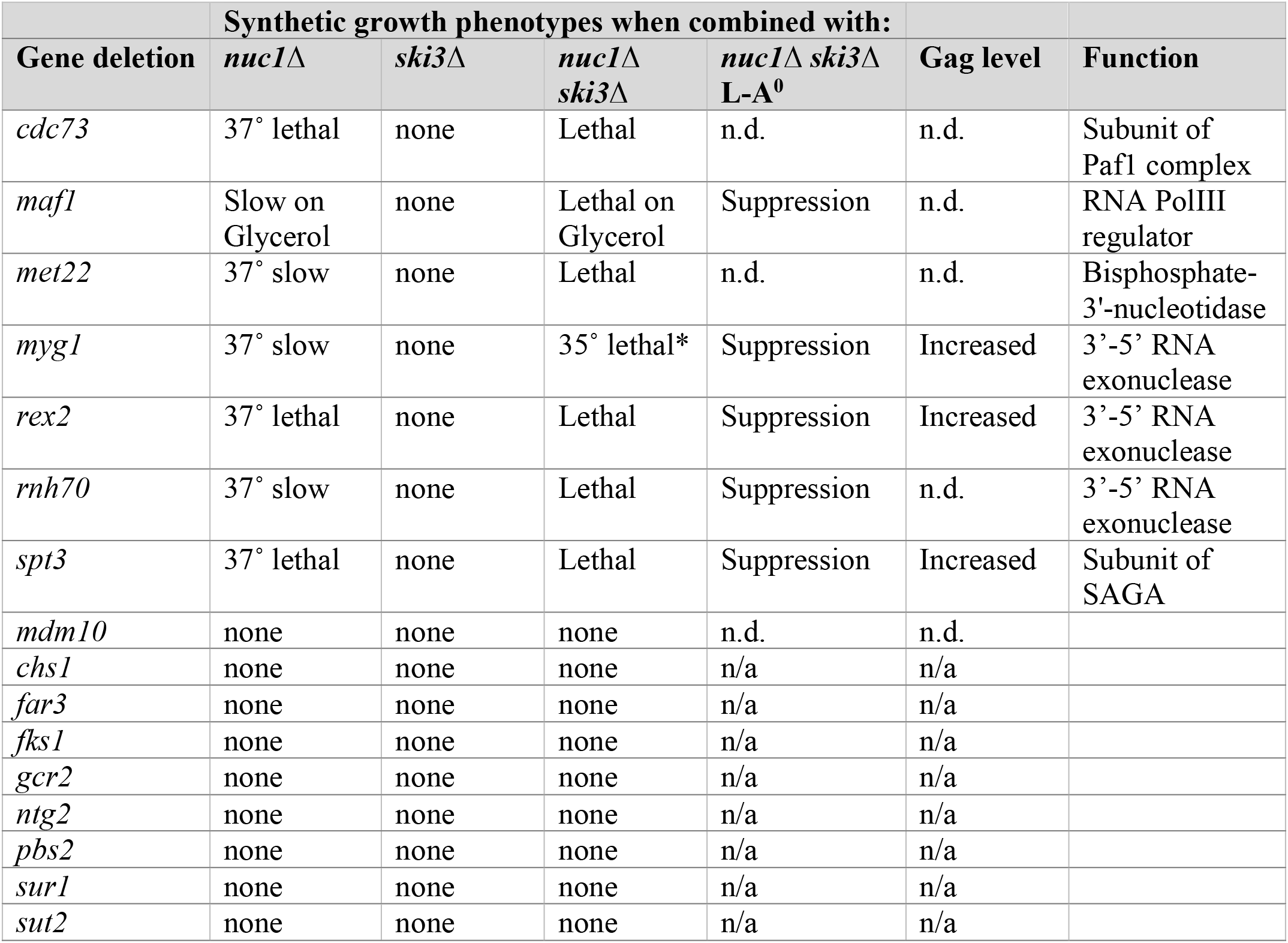
Sixteen new candidate antiviral factors were identified using a bioinformatic approach. Deletion mutants for each of these candidates were crossed to a *nuc1*Δ *ski3*Δ mutant and dissected. Phenotypes were scored as indicated and “conditional” indicates synthetic growth defects at temperatures at 37° C and/or under glycerol conditions at 30° C and 37° C. n.d. = not determined. **\***Mutants combined with *nuc1*Δ *ski3*Δ exhibit enhanced conditional growth phenotypes at 35° C compared with *nuc1*Δ *ski3*Δ.

**Table S2:**
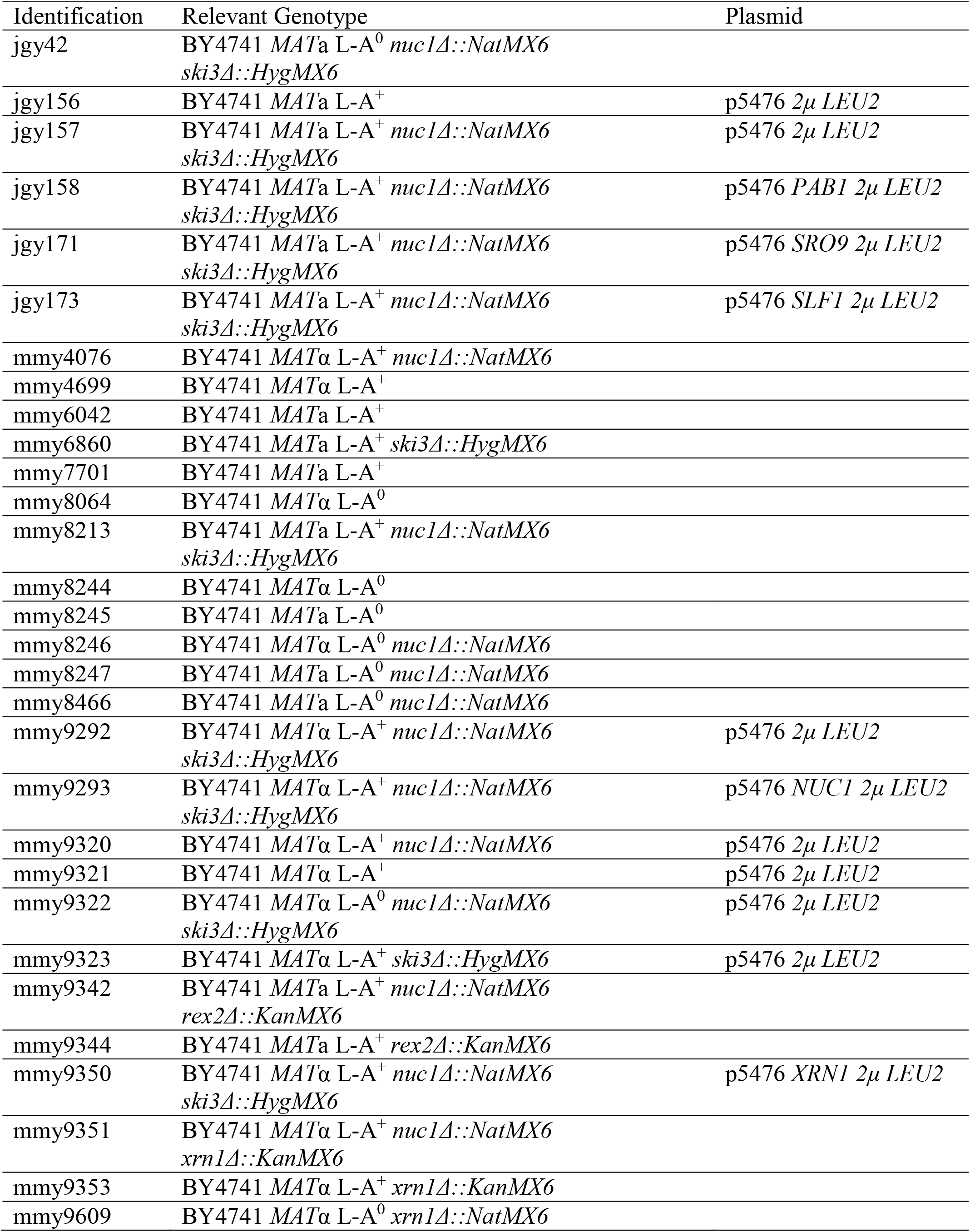
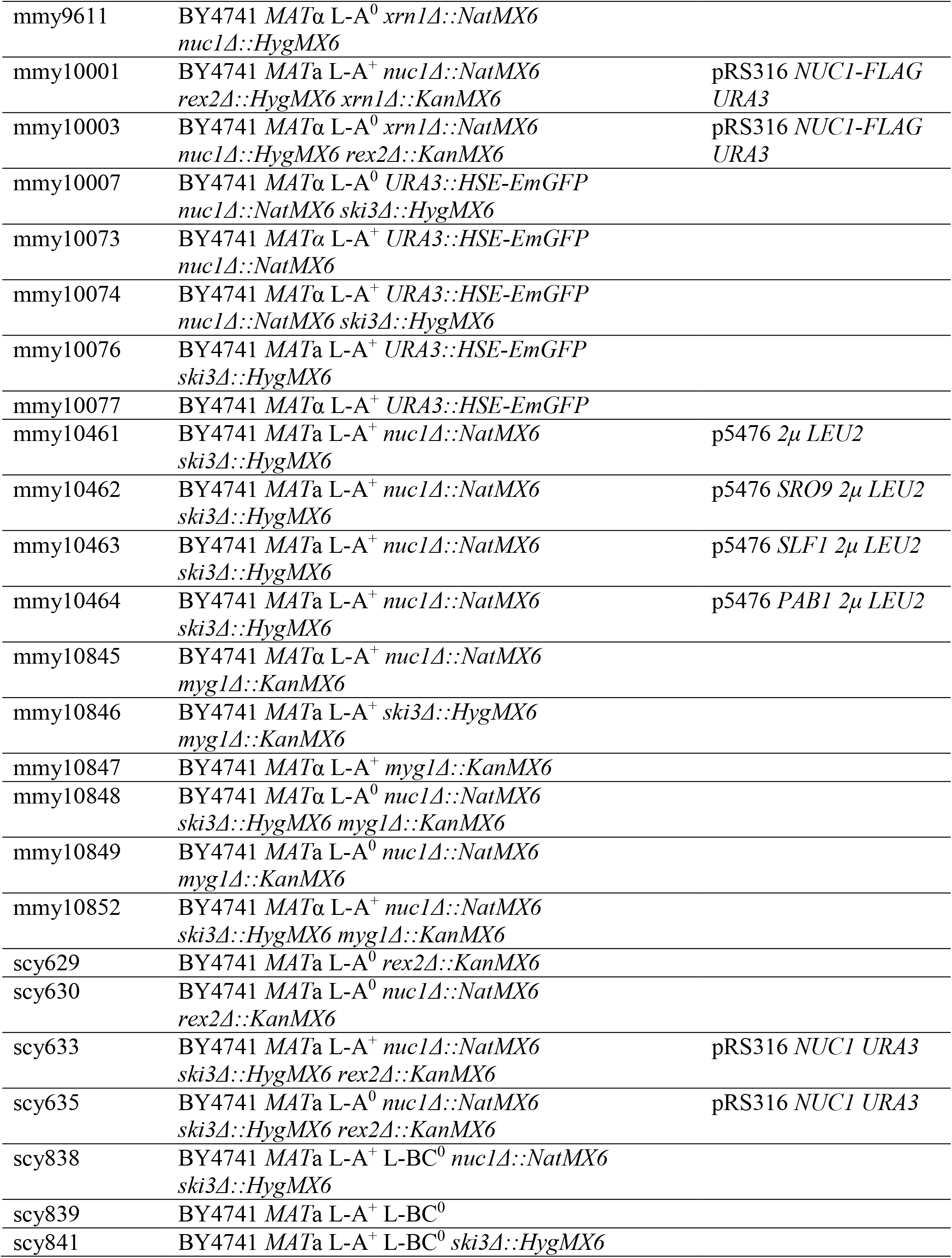
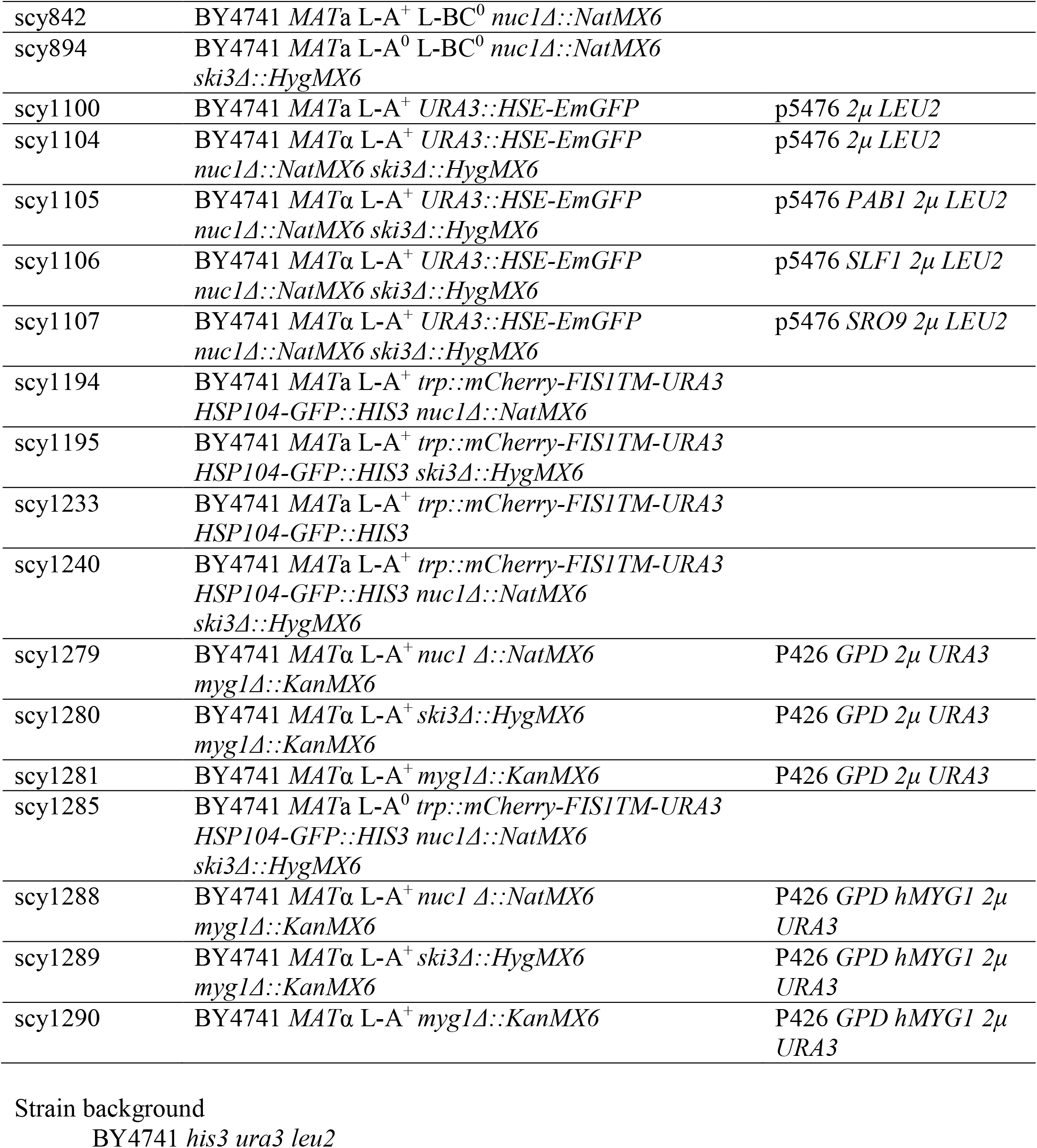
Strain and plasmid table.

